# Adaptive evolution of Topoisomerase II triggers reproductive isolation in Drosophila

**DOI:** 10.64898/2026.02.20.707007

**Authors:** Cara L. Brand, Nicholas J. Brown, Anirban Dasgupta, Michael Buszczak, Mia T. Levine

## Abstract

A fundamental driver of biological diversification is the evolution of reproductive barriers between species. Instability and mis-regulation of repetitive DNA underlie numerous post-zygotic reproductive barriers, yet the molecular mechanisms are unknown. A long-studied genetic incompatibility between *Drosophila melanogaster* and *D. simulans* arises from mis-segregation of the *D. melanogaster*-specific *359bp* DNA satellite in hybrid embryos. Here we report that the *D. simulans* version of the essential enzyme Topoisomerase II/Top2 causes this lethal incompatibility. Combining interspecies gene swaps with cell biology and genetics revealed that *D. simulans*-specific adaptive divergence of Top2 DNA-interacting domains prevents the resolution of *359bp*-induced topological stress. Our findings demonstrate that species-specific DNA satellite topology requires species-specific molecular machinery and that even vital housekeeping genes can underlie reproductive isolation between closely related species.

---

Hybrid sterility and lethality prevent gene flow between species. These post-zygotic reproductive barriers arise from deleterious interactions between species-specific DNA (Dobzhansky 1937; Muller 1942). The identity of such “speciation genes” has emerged from using classic genetic tools like recombination mapping, genome-wide screens, and introgression. These agnostic studies have revealed that the genetic loci involved in post-zygotic hybrid dysfunction support a diverse array of biological processes yet share a unifying feature: rapid sequence evolution (Orr 1995; Presgraves 2010; Frayer, et al. 2025).

The most rapidly evolving class of eukaryotic DNA sequence is repetitive DNA, which can account for up to 90% of a genome (Garrido-Ramos 2017). Extreme turnover of repetitive DNA sequence and copy number changes over short evolutionary timescales suggest that DNA repeats underlie hybrid incompatibilities. Consistently, the packaging, transmission, and integrity of DNA repeat-enriched genomic regions are frequently disrupted in hybrids (Bayes and Malik 2009; Ferree and Barbash 2009; Sanei, et al. 2011; Maheshwari, et al. 2015; Gibeaux, et al. 2018; Yoshida, et al. 2019; Jagannathan and Yamashita 2021; El Yakoubi and Akera 2023). Moreover, many speciation genes encode proteins that localize to repeat-rich genomic regions (Bayes and Malik 2009; Satyaki, et al. 2014). However, to date, there is not a single hybrid incompatibility described for which both the DNA repeat and the incompatible DNA repeat-interacting protein have been genetically identified; consequently, we have a limited grasp of the molecular mechanisms underlying these enigmatic hybrid incompatibilities. Here, we take a candidate gene approach to search directly for a DNA repeat-interacting protein responsible for a classic, DNA repeat-mediated reproductive barrier.

Genetic crosses between *Drosophila simulans* females and *D. melanogaster* males result in embryonic lethality of hybrid female offspring, but not hybrid male offspring (Fig. 1A) (Sturtevant 1920). This hybrid incompatibility maps to an *X*-linked, multi-megabase DNA satellite array called *359bp* (Sturtevant 1920; Sawamura, Yamamoto, et al. 1993; Ferree and Barbash 2009). The large, X-linked *359bp* array proliferated specifically along the *D. melanogaster* lineage and is thus absent in *D. simulans* and other closely related species (de Lima, et al. 2020). In hybrid female embryos, the paternally transmitted *359bp* satellite array causes catastrophic *X-*chromosome mis-segregation (Sawamura, Yamamoto, et al. 1993; Ferree and Barbash 2009). This incompatibility manifests early in embryogenesis when the zygotic genome is transcriptionally silent, a unique developmental stage during which maternally provisioned factors alone sustain embryonic divisions. It has long been speculated that a *D. simulans* maternally provisioned protein or RNA disrupts paternal *359bp* segregation in the hybrid embryo, but the identity of this maternal factor has remained elusive (Fig. 1A) (Sturtevant 1921; Sawamura, Taira, et al. 1993; Orr 1996; Carracedo, et al. 2000).

**Figure. 1.**
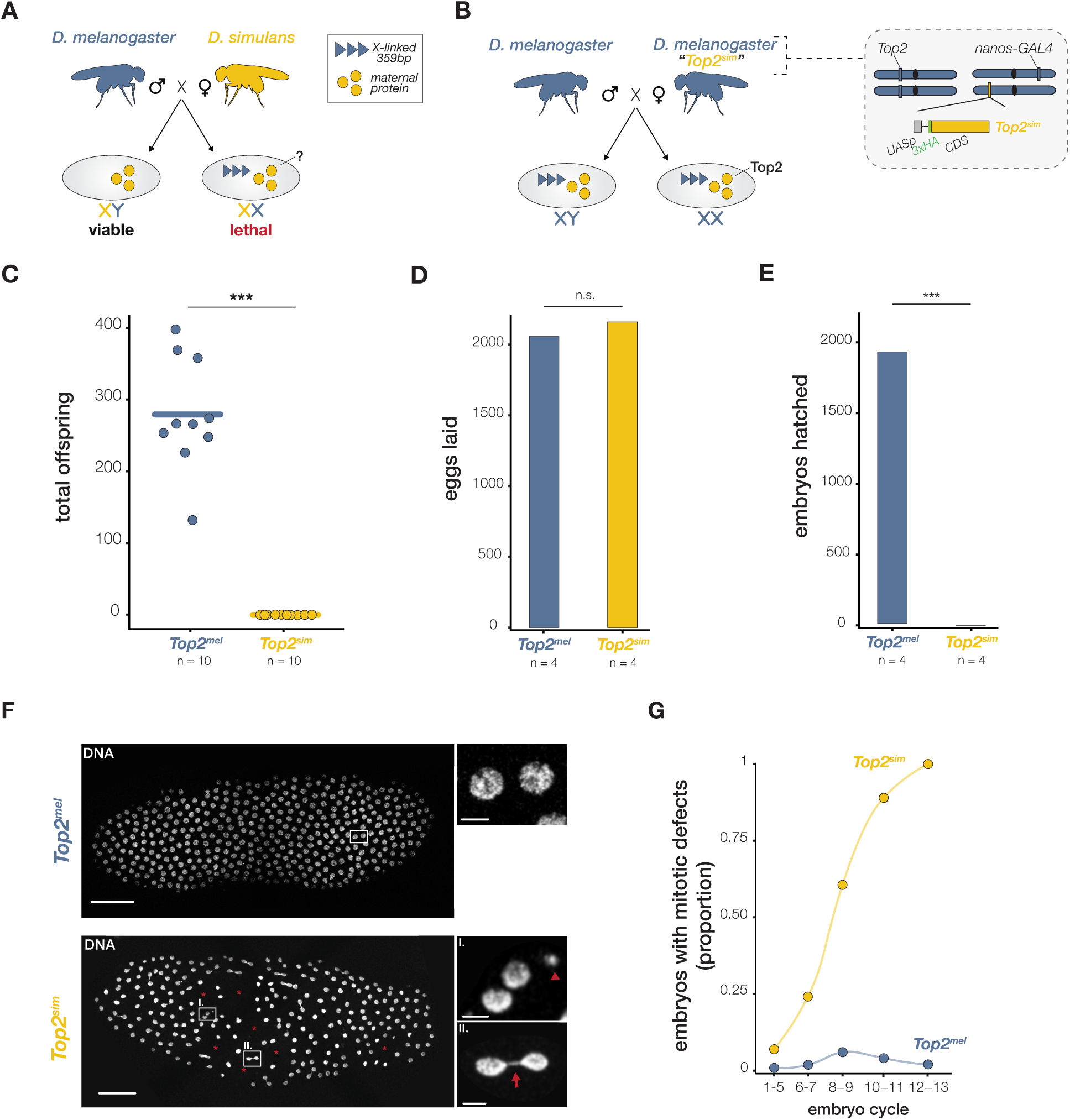
Top2 adaptive evolution results in a lethal interspecies incompatibility that manifests in the embryo. (**A**) Schematic of the early embryonic, female hybrid lethality that arises from crosses between *D. melanogaster* males and *D. simulans* females. An unknown *D. simulans* factor, maternally provisioned to the embryo, interacts deleteriously with the *D. melanogaster*-specific, *X*-linked *359bp* satellite array, causing lethality of hybrid females but not males. Hybrid males instead inherit the *D. melanogaster Y* chromosome. (**B**) Schematic of the experimental system used to assay the consequences of maternally depositing the *D. simulans* version of Top2 into *D. melanogaster* embryos. Wildtype *D. melanogaster* males are crossed to transgenic *D. melanogaster* females that encode either *D. simulans Top2* (“*Top2^sim^*”) or *D. melanogaster Top2* (“Top2^mel^”, not shown), tagged with 3xHA (green). A *nanos*-*GAL4* driver induces expression of transgenes downstream of the UASp promoter in the female germline, resulting in maternal deposition of transgenic Top2 into embryos. (**C**) Total number of offspring from *Top2^mel^* and *Top2^sim^*females crossed to wildtype males. (**D**) Number of eggs laid by *Top2^mel^*and *Top2^sim^* females. (**E**) Hatch rate of embryos from *Top2^mel^* and *Top2^sim^* females crossed to wildtype males. (**F**) Representative images of *Top2^mel^* and *Top2^sim^* embryos with nuclear fallout designated with a red asterisk (left; scale bar, 50μm). Insets display micronuclei (I, arrowhead) and chromatin bridges (II, arrow) in embryos provisioned with Top2^sim^ (scale bar, 5μm). (**G**) Quantification of embryos with mitotic defects, including chromatin bridging, micronuclei, and nuclear fallout (*n* ≥ 41 for each embryonic cycle). t-test, \*\*\**p* < 0.001

We identified the essential housekeeping enzyme Topoisomerase II (Topo II/Top2) as a strong candidate maternal factor. Top2 is maternally provisioned to *D. melanogaster* embryos (Buchenau, et al. 1993), enriched at the *359bp* array (Blattes, et al. 2006; Tang, et al. 2017), and has evolved rapidly under positive selection (Brand and Levine 2022). Across eukaryotes, Top2 maintains genome stability by altering DNA topology to relieve supercoils, resolve entanglements, and promote chromosome condensation, among other vital functions (Nitiss 2009). Top2 acts genome-wide; however, increasing evidence from flies, yeast, mouse, and human suggests that DNA repeats impose unique topological demands on Top2 function (D’Ambrosio, et al. 2008; Hughes and Hawley 2014; El Yakoubi and Akera 2023; Amoiridis, et al. 2024). We hypothesized that the torsional stress imposed by the *D. melanogaster*-specific *359bp* satellite requires the *D. melanogaster*-specific version of Top2 to maintain genome integrity. Under this model, a maternally provisioned *D. simulans* version of Top2 would be incompatible with the paternally inherited, *D. melanogaster 359bp* satellite array in the hybrid embryo.

To test this hypothesis, we first used the Gal4/UAS system to express the *D. simulans* version of Top2 in an otherwise wildtype *D. melanogaster* background. Specifically, we used the *nanos-Gal4* driver to express a 3xHA-tagged *Top2* from *D. simulans* or from *D. melanogaster* (“*Top2^sim^*” or “*Top2^mel^”*, respectively) in the *D. melanogaster* female germline. This driver leads to elevated transgene expression in the ovary and protein deposition into the egg (Fig. 1B). We found that *Top2^sim^* expression caused female sterility (Fig. 1C). Notably, these sterile females produced abundant eggs (Fig. 1D); however, these eggs failed to hatch (Fig. 1E), suggesting that maternally provisioned Top2^sim^ disrupts embryogenesis. Indeed, we observed that early embryos provisioned with Top2^sim^ (“*Top2^sim^* embryos” hereafter) exhibited severe mitotic defects, including chromosome mis-segregation and micronuclei (Fig. 1F). Such damaged nuclei are classically eliminated through a process known as nuclear fallout, which leaves large gaps within the regularly arrayed, rapidly dividing embryonic nuclei in Drosophila embryos. We observed nuclear fallout, as well as micronuclei and mis-segregating chromosomes, all of which increased in frequency across successive mitotic cycles in embryos provisioned with Top2^sim^, but not with Top2^mel^ (Fig. 1G).

To identify the genomic sequence(s) subject to mis-segregation, we performed fluorescent *in situ* hybridization using a probe specific to the *X*-linked *359bp* satellite, together with a control probe to the pericentromeric *2L3L* autosomal satellite. The *2L3L* satellite segregated to the poles in telophase nuclei in both *Top2^mel^* and *Top2^sim^* early embryos (Fig. 2A). In contrast, we observed *359bp* in the chromatin bridge of the same *Top2^sim^* early embryos (Fig. 2A), including at the first mitotic division (Fig. S1A) and at increasingly higher frequency in successive divisions, suggesting unresolved DNA torsional stress at the *X*-linked satellite array (Fig. 2B). Only in later divisions did *2L3L* appear in the bridge, and always coincident with *359bp*. These findings demonstrate that *D. melanogaster* maternal provisioning of Top2^sim^ triggers *359bp* mis-segregation in early embryonic divisions, followed by global genome instability in later cycles.

**Figure. 2.**
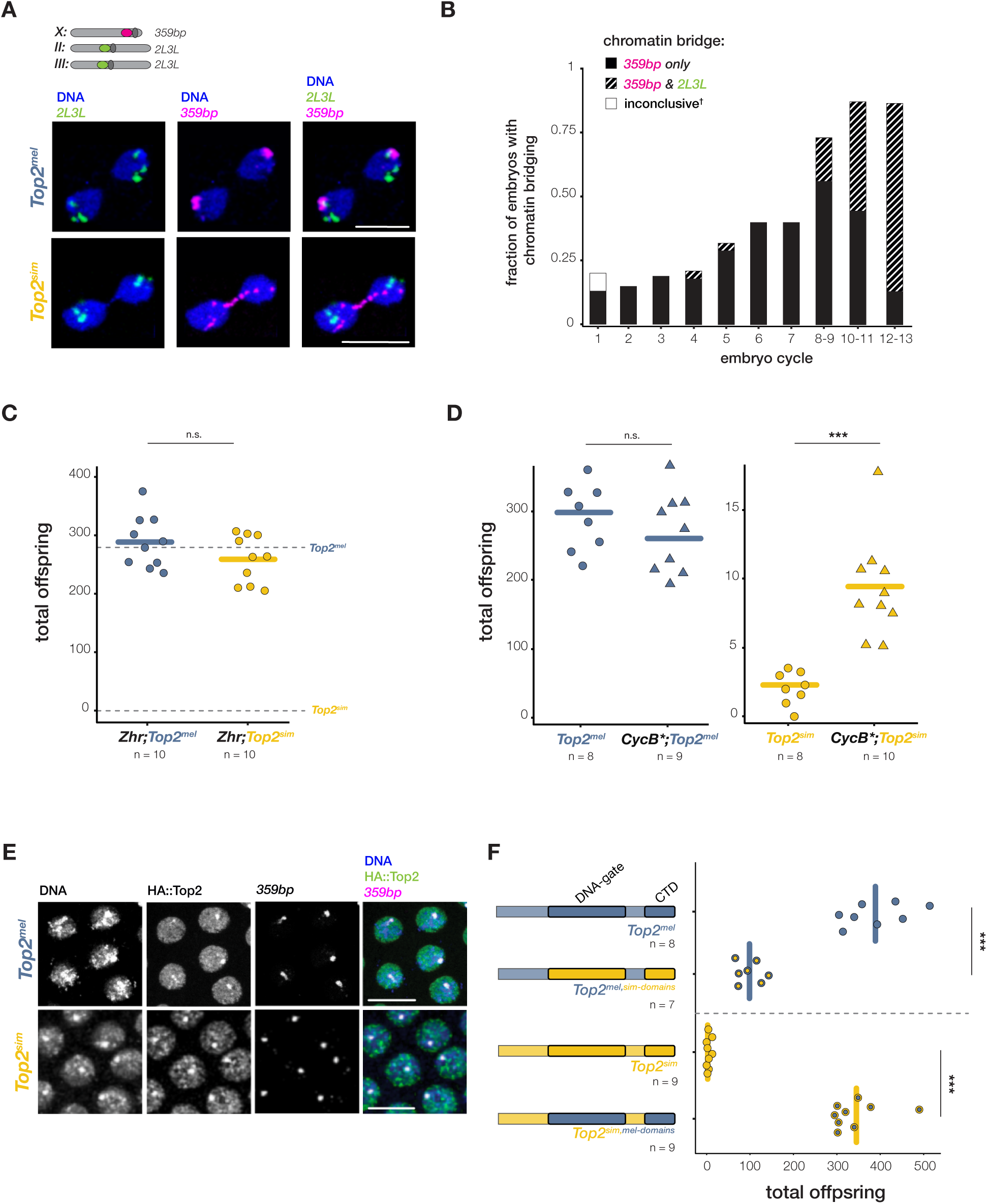
Maternal deposition of the *D. simulans* Top2 causes *359bp* mis-segregation in the *D. melanogaster* early embryo. (**A**) Schematic of the genomic locations where FISH probes anneal and representative images of *2L3L* (green) and *359bp* (pink) at telophase in Top2^mel^- and Top2^sim^-provisioned syncytial embryos (scale bar, 10μm). (**B**) Frequency of chromatin bridging observed in Top2^sim^-provisioned embryos and proportion of embryos with bridges containing *359bp* alone, or *359bp* and *2L3L* (n ≥ 8 for each embryonic cycle). ^†^We observed a single mis-segregation event with neither probe detected in the bridge; however, the positioning and shape of the *359bp* signal were aberrant (fig. S1B). (**C**) Total offspring from *Zhr;Top2^mel^* and *Zhr;Top2^sim^*females lacking the *X*-linked *359bp* array. Mean offspring of *Top2^mel^* and *Top2^sim^* females from Fig. 1C are marked with dashed lines. (**D**) Total offspring from *Top2^mel^* and *CycB*;Top2^mel^* females (left) and *Top2^sim^* and *CycB*;Top2^sim^* females (right). (**E**) Representative images of HA::Top2 (green) and *359bp* (pink) colocalization in *Top2^mel^* and *Top2^sim^* syncytial embryos (scale bar, 10μm). (**F**) Total offspring from females expressing chimeric Top2 constructs. The adaptively evolving DNA-gate and C-terminal domains are outlined in black. CTD = C-terminal domain. t-test, ***p < 0.001.

To test whether the observed *359bp* chromatin bridging in Top2^sim^-provisioned embryos caused lethality, we took advantage of the *Zygotic hybrid rescue* (*Zhr*) mutant, which lacks the *X*-linked, 11Mb *359bp* satellite array (Sawamura, Yamamoto, et al. 1993). Specifically, we crossed *Zhr* males to females expressing Top2^sim^ in a *Zhr* genetic background. The resulting embryos are provisioned with Top2^sim^ but lack the *359bp* satellite array. The *359bp* deletion fully rescued embryo viability (Fig. 2C), demonstrating that embryonic lethality arises from a deleterious interaction between Top2^sim^ and the *359bp* satellite. This finding reveals a new genetic incompatibility that arises from a deleterious interaction between a *D. simulans*-specific allele and a *D. melanogaster*-specific satellite.

Maternal provisioning of Top2^sim^ triggered catastrophic *359bp* mis-segregation during embryogenesis; however, Top2^sim^ overexpression in the female germline had no detectable impact on oogenesis (Fig. 1D, S1C,D), implicating an embryo-specific vulnerability. Indeed, we detected no evidence of lethality upon ubiquitously expressing Top2^sim^ after zygotic genome activation in the embryo, or in the larval and adult stages (Fig. S1E). Unlike other developmental stages, the early embryo undergoes extremely rapid mitotic divisions (Foe and Alberts 1983).

We hypothesized that this rapid cycling underlies the observed embryo-specific susceptibility. Under this model, slowing embryonic cell cycles would alleviate the mitotic defects triggered by Top2^sim^. To test this prediction, we reduced the maternal dose of a key cell cycle regulator, Cyclin B, resulting in slightly lengthened embryonic cell cycles (Ji, et al. 2004). This very modest increase in cell cycle length was sufficient to increase embryonic viability (Fig. 2D). These data suggest that, despite the presence of endogenous *D. melanogaster* Top2 (Fig. 1B), Top2^sim^ disrupts the resolution of torsional stress imposed by the *359bp* array during the rapid nuclear divisions. More specifically, Top2^sim^ likely recognizes *359bp* but has insufficient time to promote faithful *X* chromosome segregation. Consistent with this model, we found that Top2^sim^, like Top2^mel^, is enriched at *359bp* in *D. melanogaster* embryos (Fig. 2E) and Top2^sim^ hyperaccumulates on the satellite array relative to transgenic Top2^mel^ in ovaries, a tissue where Top2^sim^ expression is benign (see Materials and Methods, Fig. S2). These data suggest that while Top2^sim^ can recognize this problematic locus, adaptive evolution altered its efficiency in resolving satellite-induced topological stress in the rapidly cycling embryo.

To explore the molecular basis of Top2^sim^ inefficiency at *359bp*, we analyzed adaptive sequence evolution across the distinct functional domains of Top2. These domains coordinate the resolution of torsional stress through an ATP-dependent strand-passage reaction (Nitiss 2009). The “DNA-gate” covalently cross-links to one DNA strand, creating a transient double-strand break. Two other structural domains – the “N-gate/ATPase domain” and “C-gate” – promote the capture and passage of a second DNA strand. The DNA-gate then re-ligates the double-strand break. The C-terminal domain mediates Top2 interactions with DNA and chromatin proteins, regulating enzyme recruitment, processivity, and efficiency (Clarke and Azuma 2017). A sliding window McDonald–Kreitman test (McDonald and Kreitman 1991) across the Top2 coding region revealed an excess of nonsynonymous fixations within the DNA-gate (Fig. S3A), and a sliding window dN/dS analysis uncovered elevated nonsynonymous divergence across the C-terminal domain (Fig. S3B). Such signatures of positive selection are consistent with a history of domain-specific adaptation. These two DNA-interacting regions of Top2, in contrast to its two structural domains, modulate the efficiency of strand passage (Froelich-Ammon, et al. 1994; Lane, et al. 2013; Antoniou-Kourounioti, et al. 2019; Jang, et al. 2019), suggesting that adaptive divergence of the DNA-gate and the C-terminal domain underlies species-specific differences in Top2 function at the *D. melanogaster*-specific *359bp*.

To determine whether adaptive amino acid changes across the DNA-gate and C-terminal domain impact Top2 function, we assayed the fertility of females that maternally provision embryos with chimeric Top2 proteins. *D. melanogaster* females expressing a chimeric Top2^mel^ with the *D. simulans* DNA-gate and C-terminal domains were severely subfertile, while expression of a chimeric Top2^sim^ with the *D. melanogaster* DNA-gate and C-terminal domains fully rescued female fertility (Fig. 2F). Swapping either the *D. melanogaster* DNA-gate alone or C-terminal domain alone into Top2^sim^ revealed that each domain contributed to the rescue; however, these domains act synergistically such that the rescue by the chimera with both *D. melanogaster* domains exceeded additivity (Fig. S3C). This series of chimeras, combined with the cell cycle length-dependent rescue above (Fig. 2D), suggests that *D. melanogaster*-specific residues within both DNA-interacting domains are required to efficiently execute the strand-passage reaction at *359bp*.

The characterized functions of the DNA-gate and C-terminal domain offer mechanistic insight into how adaptive divergence may alter Top2 activity at *359bp*. Secondary structures enriched at satellites alter how the DNA-gate engages and bends DNA (Howard, et al. 1991; Burden and Osheroff 1999; Jang, et al. 2019), impacting Top2 cleavage efficiency (Froelich-Ammon, et al. 1994; Szlachta, et al. 2020). We speculate that *D. simulans*-specific amino acid changes within the DNA-gate restrict Top2-dependent bending and cleavage of satellite-specific structures formed at the *359bp* array (Borgnetto, et al. 1999). Moreover, secondary structure-prone satellite DNA introduces topological complexity that requires repeated rounds of strand passage (Lane, et al. 2013; Lee, et al. 2023). Previous work has shown that the Top2 C-terminal domain is required to meet these demands by promoting efficient Top2 activity through interactions with DNA as well as local histones and non-histone chromatin-associated factors (Lane, et al. 2013; Antoniou-Kourounioti, et al. 2019; Shintomi and Hirano 2021). We speculate that *D. simulans*-specific residues across the Top2^sim^ C-terminal domain fail to promote these local interactions. Reduced functional engagement at *359bp*, together with reduced bending of the *359bp* sequence, may underlie the observed Top2^sim^ inefficiency and consequent topological stress that triggers lethal mis-segregation in *D. melanogaster* embryos.

Maternal provisioning of transgenic Top2^sim^ to *D. melanogaster* embryos leads to defects that closely resemble those observed in hybrid female embryos produced from crosses between wild-type *D. simulans* females and *D. melanogaster* males (Fig. 1A). This striking parallel implicates *D. simulans* Top2 as the elusive maternal factor incompatible with the *D. melanogaster*-specific *359bp* satellite. Under this model, providing *D. melanogaster* Top2 to hybrid embryos should rescue hybrid female viability. To test this, we expressed Top2^mel^ in *D. simulans* under the maternal ɑ-tubulin promoter, which drives expression in late oogenesis and leads to maternal protein deposition into the egg (Fig. 3A) (Matthews, et al. 1989). Control crosses between wildtype *D. melanogaster* males and wildtype *D. simulans* females resulted in nearly 100% penetrant hybrid female lethality. Experimental crosses between wildtype *D. melanogaster* males and the *D. simulans* females expressing Top2^mel^ resulted in 10-fold more female progeny (Fig. 3B). These data indicate that *D. simulans* Top2 underlies the F1 hybrid incompatibility with *D. melanogaster 359bp*, a reproductive barrier reported over 100 years ago.

**Fig. 3.**
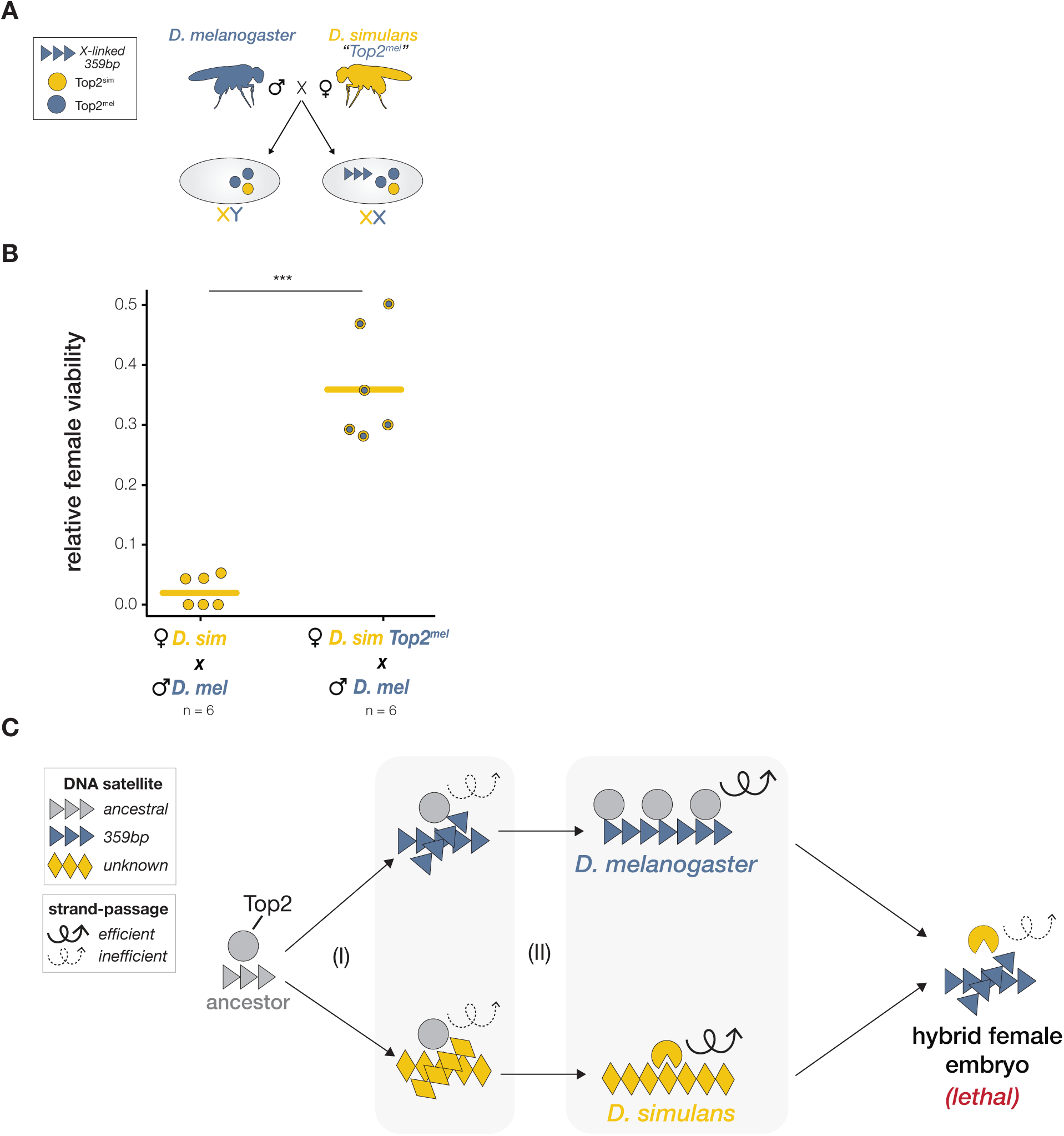
An incompatibility between Top2 and *359bp* causes an F1 reproductive barrier between *D. melanogaster* and *D. simulans*. (**A**) Schematic of the experimental system used to test hybrid rescue. Wildtype *D. melanogaster* males are crossed to transgenic *D. simulans* females (strain MD199S, see Materials and Methods) expressing Top2^mel^ under a maternal-α tubulin promoter, leading to maternal deposition of transgenic Top2^mel^ into hybrid embryos. (**B**) Relative female offspring viability from crosses between wildtype *D. melanogaster* (“*D. mel*”) males and wildtype *D. simulans* females (“*D. sim*”) and between wildtype *D. melanogaster* males and transgenic *D. simulans* females expressing Top2^mel^ (“*D. sim Top2^mel^*”). t-test, ***p < 0.001. (**C**) Model of the causes and consequences of Top2 evolution. (I) Lineage-specific evolution of satellite DNA leads to novel topological stress that reduces the efficiency of the ancestral Top2 strand passage reaction. (II) A fitness reduction leads to selection for altered Top2 abundance or amino acid sequence. Along the *D. melanogaster* lineage (upper path), Top2 evolves increased abundance through reduced activity of the adaptively evolving protease, Maternal Haploid (Brand and Levine 2022) (not pictured), which targets Top2 for degradation. Along the *D. simulans* lineage (lower path), Top2 itself evolves adaptively to engage novel secondary structures and alter residence time at an unknown DNA satellite. Both evolutionary trajectories restore efficient strand-passage. The adaptive divergence of *D. simulans* Top2 is incompatible with the topological demands of the *D. melanogaster 359bp* satellite array, triggering embryonic lethality of hybrid females.

The penetrance of maternal-effect hybrid lethality varies continuously across *D. simulans* strains, ranging from complete embryonic lethality to complete embryonic viability of hybrid daughters (Gerard and Presgraves 2012). This quantitative variation suggests that, unlike most hybrid incompatibility factors, this *D. simulans* maternal-effect hybrid lethality is multigenic (Gerard and Presgraves 2012). Under this framework, variation at Top2, along with one or more additional loci, contributes to female hybrid lethality. We found that five focal *D. simulans* strains with a wide range of maternal-effect lethality (fig. S4) share identical *Top2* amino acid sequences (Fig. S5), suggesting that protein sequence variation at *Top2* does not contribute to the observed phenotypic variation. We hypothesize that variation at other loci across the genome determines penetrance of Top2-mediated hybrid lethality, possibly by modulating maternal protein deposition.

Our observation that a fast-evolving DNA repeat from one species interacts deleteriously with a fast-evolving DNA repeat-interacting protein from another suggests a classic model of rapid, lineage-specific DNA-protein coevolution (Henikoff, et al. 2001; Presgraves 2010; Maheshwari and Barbash 2011; Ferree and Prasad 2012). Under this model, rapid *359bp* proliferation along the *D. melanogaster* lineage triggered Top2 adaptation to maintain efficient strand-passage activity. However, we uncovered evidence that Top2 accumulated adaptive changes along the *D. simulans* branch, not the *D. melanogaster* branch (fig. S3D). This unexpected result suggests that Top2 adaptation was not shaped by *359bp* but instead by coevolution with a *D. simulans*-specific genetic element, possibly a *D. simulans*-specific DNA satellite, that rendered Top2^sim^ inefficient at the *359bp* array (Fig. 3C). Intriguingly, previous research suggests that *359bp* proliferation selected for increased Top2 abundance rather than changes to the protein sequence (Fig. 3C). Maternal Haploid/SPARTN is a protease that targets Top2 for degradation (Lopez-Mosqueda, et al. 2016). *D. melanogaster* Maternal Haploid (Loppin, et al. 2001; Tang, et al. 2017) appears to have adaptively evolved reduced protease activity, increasing Top2 abundance to meet the topological demands of *359bp* (Brand and Levine 2022). The two studies, in combination, reveal that the proliferation of a single satellite array can generate multiple incompatibilities involving distinct members of a single pathway.

The observation that a housekeeping gene, shared across eukaryotes, mediates an interspecies incompatibility was unexpected. However, we discovered that signatures of Top2 adaptive evolution are not restricted to the *D. simulans* lineage. Top2 evolves adaptively across rodent and primate phylogenies as well as across multiple Drosophila lineages (fig. S6). These molecular evolution data suggest that incompatibilities between Top2 and DNA repeats may be widely distributed. Indeed, virtually all eukaryotic genomes contain vast stretches of rapidly evolving DNA satellites that can impose extreme topological stress, threatening genome integrity (D’Ambrosio, et al. 2008; Hughes and Hawley 2014; El Yakoubi and Akera 2023; Amoiridis, et al. 2024).

Historically, most DNA repeats were omitted from genome assemblies due to various technical barriers and a lack of interest in these ostensibly non-functional parts of the genome. The recent surge of telomere-to-telomere genome assemblies of rodents, primates, insects, plants, and other eukaryotes has remedied this omission (Rhie, et al. 2021; Nurk, et al. 2022; Chen, et al. 2023; Francis, et al. 2025; Yoo, et al. 2025; Zhang, et al. 2025). Comparisons of long-read assemblies across closely related species (Kim, et al. 2021; Rhie, et al. 2021; Logsdon, et al. 2024; Yoo, et al. 2025) have revealed a striking diversity of fast-evolving repetitive sequences that likely impose lineage-specific topological challenges. We anticipate that integrating these repeat-rich genomic regions with evolution-guided functional analyses will reveal how dynamic repetitive landscapes reshape even the most conserved biological pathways and erect reproductive barriers between species.

## Materials and Methods

### Molecular evolution analyses

To identify Top2 domains enriched for positive selection, we performed sliding window McDonald–Kreitman tests (McDonald and Kreitman 1991) on nine *D. melanogaster Top2* alleles (coordinates 2L:19447365-19453490, dmel r6.4) from lines collected in Lyon, France (Pool, et al. 2012), and seven *D. simulans Top2* alleles (Brand and Levine 2022) from lines collected in Nairobi, Kenya (Rogers, et al. 2014) (data file S1). We generated a multiple sequence alignment with the Geneious Alignment algorithm using default settings (Geneious Prime, Biomatters, Auckland, New Zealand) and visually confirmed alignment quality. We performed sliding window McDonald–Kreitman tests across the Top2 coding region (window size = 200 codons, step size = 20 codons). We also calculated pairwise dN/dS between *D. melanogaster* and *D. simulans* alleles across the Top2 coding region (window size = 100bp, step size = 20bp) (Librado and Rozas 2009). To assay lineage-specific evolution, we performed McDonald–Kreitman tests (McDonald and Kreitman 1991) using the *D. yakuba Top2* coding sequence as an outgroup to polarize mutations along the *D. melanogaster* and *D. simulans* lineages.

To test for evidence of recurrent adaptive evolution, we conducted a phylogeny-based molecular evolution analysis of *Top2* orthologs across 17 species within the *melanogaster* group (Fig. S6A). We generated a multiple sequence alignment with the Geneious Alignment algorithm using default settings (Geneious v11.1.5, Biomatters, Auckland, New Zealand). We visually confirmed alignment quality and gap-adjusted the alignment to retain in-frame codons. We fit the multiple sequence alignment to NSsites models using the codeml package in PAML (Yang 1997). To determine statistical significance, we performed a likelihood ratio test, comparing model 7 (dN/dS values fit a beta distribution between 0 to 1) and model 8 (model 7 parameters plus an additional class allowing dN/dS > 1). We also analyzed Top2 molecular evolution across 15 rodent species (Fig. S6B) and 15 primate species (Fig. S6C) using the codeml package in PAML within the FREEDA pipeline (Dudka, et al. 2023). We performed separate analyses for the two Top2 paralogs, Top2A and Top2B, present in mammals.

### Fly stock construction

#### Constructing transgenic Top2 overexpression lines with and without other mutations

We used the *ΦC31* integrase-mediated transgenesis system (Venken, et al. 2006; Bischof, et al. 2007) to insert *Top2* alleles from *D. melanogaster* (Brand and Levine 2022) or *D. simulans* downstream of a UAS promoter into the same genomic landing site. We synthesized codon-optimized *Top2* coding sequences from both species (Twist Bioscience, South San Francisco, CA), adding an N-terminal 3xHA tag and linker sequence (GGTGGTTCATCA). We cloned the resulting constructs into the *NotI*/*SpeI* sites of the pUASp-attB vector (Drosophila Genomics Resource Center, Bloomington, IN). We injected the constructs into *D. melanogaster yw; PBac[y⁺-attP-9A]VK00018* flies, which carry an attP landing site at cytological position 75A10 on chromosome *3L* (BestGene, Chino Hills, CA). We generated homozygous *UASp-Top2^mel^* or *UASp-Top2^sim^*transgenic lines using balancer chromosomes. To drive expression of the transgenic *Top2* alleles in the female germline, we crossed the lines to *nos-Gal4::VP16* (BDSC #64277). To drive ubiquitous expression, we crossed the lines to *Act5C-Gal4* (BDSC #3954).

We also used the *ΦC31* integrase-mediated transgenesis system to introduce chimeric *D. melanogaster-D. simulans Top2* alleles downstream of a UAS promoter. We constructed four chimeric Top2 constructs, each encoding an N-terminal 3xHA tag and linker sequence (GGTGGTTCATCA) and cloned them into the *NotI/SpeI* sites of the pUASp-attB vector (Drosophila Genomics Resource Center, Bloomington, IN). Specifically, we engineered *D. melanogaster Top2* alleles that code for both the *D. simulans* DNA-gate (amino acids 410-1041) and C-terminal domain (amino acids 1200–1446). We also constructed the reciprocal *D. simulans Top2* alleles encoding both the *D. melanogaster* DNA gate and C-terminal domain. Finally, we engineered two *D. simulans Top2* alleles that code for either the *D. melanogaster* DNA-gate only or the C-terminal domain only. We introduced these constructs into *D. melanogaster yw; PBac[y^+^-attP-9A]VK00018* flies (see above, BestGene, Chino Hills, CA). We drove expression of the chimeric Top2 transgenes in the female germline using *nos-Gal4::VP16* (BDSC #64277).

We used a *+/FM7; +/TM6* stock to construct flies that contain both the *359bp* satellite deletion (*Zhr*^1^, BDSC #25140) on the *X* chromosome and either the *UASp-Top2^mel^* or *UASp-Top2^sim^* transgene on chromosome *3*. Similarly, we used the *+/FM7; +/TM6* stock to construct flies carrying both the *359bp* satellite deletion (*Zhr^1^*, BDSC #25140) on the *X* chromosome and the *nos-Gal4::VP16* driver (BDSC #64277) on chromosome *3*. We crossed either *Zhr;UASp*-*Top2^mel^*or *Zhr;UASp-Top2^sim^* females to *Zhr;nos-Gal4::VP16* males to generate females that overexpress each transgene in a *359bp* deletion background.

We used a *+/FM7; +/CyO* stock to construct flies that contain a heterozygous *CycB^2^/CyO* mutation (Jacobs, et al. 1998) on chromosome *2* and either the *UASp-Top2^mel^* or *UASp-Top2^sim^* transgene on chromosome *3*. We crossed *nos-Gal4::VP16* (BDSC #64277) females to either *CycB^2^/CyO; UASp-Top2^mel^* or *CycB^2^/CyO; UASp-Top2^sim^*males to generate females that overexpress each transgene in either a *CyO* or *CycB^2^* hemizygous background. Note that we attribute the additional survivors from *CyO;Top2^sim^* females (see Fig. 1D) to a distinct genetic background from *Top2^sim^* females (see Fig. 1C).

#### Constructing a transgenic D. simulans line expressing Top2^mel^

We used CRISPR/Cas9 to generate *D. simulans* flies carrying a transgenic *D. melanogaster* allele of *Top2*. We modified an existing Janelia Atalanta plasmid (pJAT17; (Stern, et al. 2023)) to encode a codon-optimized *D. melanogaster Top2* coding sequence driven by the maternal-*α* tubulin promoter and 3’ UTR, as well as a 3xP3 driven EYFP marker. This construct was designed for site-specific insertion at 3R: 9,189,697 using a U6 promoter-driven guide RNA (GCACACACGCCCATACAAAG) and homology arms targeting ∼600bp upstream and downstream of the gRNA target site. The donor vector was co-injected with Cas9 mRNA into *D. simulans* strain MD199S (SKU:1421-0241.197; Rainbow Transgenics, Camarillo, CA).

### Fertility, fecundity, and embryo viability assays

We performed all experiments at least twice (N ≥ 2 except where noted).

We conducted all experiments on standard cornmeal food at 25°C, except where noted. To assay female fertility, we aged virgin females for three to five days. For each replicate vial, we crossed four virgin females with four males. We transferred parents to fresh vials every three days over a nine-day period and counted all progeny that emerged.

To assay female fecundity and embryo viability, we allowed ∼200 mated females to lay eggs on Nutri-fly grape agar media (Genesee Scientific, El Cajon, CA) for one hour after a one-hour pre-lay. We then counted the total number of embryos laid and the number of larvae that hatched each day over a 72-hour period.

To assay adult viability, we crossed four virgin *UASp-Top2^mel^ or UASp-Top2^sim^* females to *Act5C-GAL4/TM6* (BDSC #3954) males (N ≥ 1). We transferred parents to fresh vials every three days over a nine-day period and counted all progeny that emerged. We calculated viability as the proportion of *UAS/GAL4* progeny relative to their *UAS/TM6* balancer siblings from the same cross.

To assay hybrid embryo viability, we crossed 30 *D. simulans* virgin females with 50 *D. melanogaster males*. We transferred parents to fresh vials every three days until all females had died. For each interspecies cross, we analyzed the sex ratio only of replicates that produced at least 25 progeny. We performed these crosses under constant light at 18°C, as hybrid female viability is higher at lower temperatures (Gerard and Presgraves 2012).

### Embryo collection and cytology

After a 1-hour pre-lay, we collected embryos on grape juice agar plates, dechorionated in 50% bleach, and fixed and devitellinized in 100% methanol and heptane (Rothwell and Sullivan 2007).

We conducted fluorescence *in situ* hybridization following the protocol described in (Dernburg 2011). We designed fluorophore-conjugated probes (Integrated DNA Technologies, Coralville, IA) of the *359bp* satellite array (5’Alex594N:TTTTCCAAATTTCGGTCATCAAATAATCAT) on the *X* chromosome and the *2L3L* satellite arrays (5’Alex488N: AATAACATAG3) on the second and third chromosomes. We co-hybridized the probes at 32°C overnight and mounted embryos with ProLong Gold Antifade Reagent with DAPI (Thermo Fisher Scientific, Waltham, MA). We imaged slides at 63x magnification on a Leica TCS SP8 Four Channel Spectral Confocal System.

We conducted immunofluorescence with rat anti-HA antibody clone 3F10 (1:1000; Cat#: ROAHAHA, RRID: AB_2687407, Sigma-Aldrich, St. Louis, MO) and a goat anti-rat fluorophore-conjugated secondary antibody (1:500; Cat# A-11007, RRID: AB_141374, Thermo Fisher Scientific, Waltham, MA) following the protocol described in (Loppin, et al. 2001). We mounted and imaged the embryos as described above.

We performed all embryo collection and staining experiments at least twice (N ≥ 2).

### ChIP-qPCR

To assay Top2 enrichment at the *359bp* array, we performed chromatin immunoprecipitation followed by quantitative PCR (ChIP-qPCR). We performed ChIP-qPCR in ovaries to measure baseline Top2 enrichment at the *359bp* array, rather than secondary recruitment arising from segregation defects in embryos. For each biological replicate (N = 3), we dissected 50 pairs of ovaries from *Top2^mel^* and *Top2^sim^* females (Fig. 1B). We crosslinked ovaries in 1mL A1 buffer (10mM sodium phosphate dibasic, 2mM potassium phosphate monobasic, pH 7.4, 137mM sodium chloride, 2.7mM potassium chloride, 1% BSA, and cOmplete EDTA-free Protease Inhibitor Cocktail (Roche, Basel, Switzerland)) supplemented with 1% formaldehyde for 15 minutes at room temperature. We quenched crosslinking by adding 125mM glycine for 5 minutes, then washed ovaries in ice-cold A1 buffer.

To prepare fragmented chromatin for immunoprecipitation, we homogenized and lysed ovaries in lysis buffer (10mM Tris-HCl pH 8.0, 140mM sodium chloride, 1mM EDTA, 1% Triton X-100, 0.1% SDS, 0.1% sodium deoxycholate, and protease inhibitors) on a rotator at 4°C for 1 hour. We sonicated chromatin using a Covaris E220 for 15 minutes (30s on/30s off, high power), yielding fragments of 400-500bp, confirmed by agarose gel electrophoresis.

To immunoprecipitate Top2-associated chromatin, we reserved 5% input from the supernatant and precleared the remaining chromatin with 20uL Dynabeads Protein A (Thermo Fisher Scientific, Waltham, MA). We incubated precleared chromatin overnight at 4°C with 5ug rabbit anti-HA antibody (Cat# ab9110, RRID: AB_307019, Abcam, Cambridge, UK) bound to 50uL Dynabeads Protein A. We collected HA::Top2-bound chromatin by centrifugation and washed beads using standard low-salt, high-salt, LiCl, and TE buffers. To purify DNA for downstream qPCR analysis, we eluted chromatin in 50mM Tris-HCl (pH 8.0), 10mM EDTA, 1% SDS at 65°C for 15 minutes. We incubated the eluate overnight at 65°C to reverse crosslinks and treated samples with RNase A and Proteinase K (Thermo Fisher Scientific, Waltham, MA). We purified DNA using the Zymo DNA Clean & Concentrator Kit (Zymo Research, Irvine, CA) and eluted in nuclease-free water.

For each biological replicate, we performed three qPCR technical replicates using primers targeting either the *359bp* array (F: GTTTTGAGCAGCTAATTACC and R: TATTCTTACATCTATGTGACC) or the telomeric retrotransposon HeT-A (F: CGCGCGGAACCCATCTTCAGA and R: CGCCGCAGTCGTTTGGTGAGT) as a negative control. We calculated enrichment as percent input relative to the corresponding input sample.

### Natural variation of Top2 across *D. simulans*

To quantify natural variation across the *D. simulans* Top2 coding sequence, we designed primers that amplified the 4.5kb Top2 gene region from five strains (NCBI BioSample: sim6: 14021-0251.194, w501: 14021-0251.195, MD199S: 14021-0251.197, NC48S: 14021-0251.198, sim4: 4021-0251.216). We then prepared genomic DNA and conducted PCR amplification followed by Sanger sequencing using standard protocols. We aligned the sequences in Geneious using the Geneious Alignment algorithm with default settings (Geneious Prime, Biomatters, Auckland, New Zealand) and confirmed alignment quality by eye.

## Acknowledgments

We thank the Levine Lab, Lampson Lab, P. Geyer, D. Presgraves, D. Dudka, and A. Ridgway for feedback on the manuscript. We also thank S. Mohammed for technical assistance and E. Behrman and E. Marti for their expertise and guidance on the construction of the transgenic *D. simulans* fly and D. Presgraves for generously sharing fly strains. This work was supported by the National Institutes of Health grants K99GM149943 to CLB, T32GM153602 to NJB, R35GM144043 and R01AG079513 to MB, and R35GM124684 to MTL and the Welch Foundation grant I-2226-20250403 to MB.

## Author contributions

Conceptualization: CLB, MTL, Investigation: CLB, NJB, AD, Supervision: MTL, MB, Visualization: CLB, MTL, Writing – original draft: CLB, MTL , Writing – review & editing: CLB, NJB, AD, MB, MTL

