## Supplementary Material for "Adaptive evolution of Topoisomerase II triggers reproductive isolation in Drosophila"

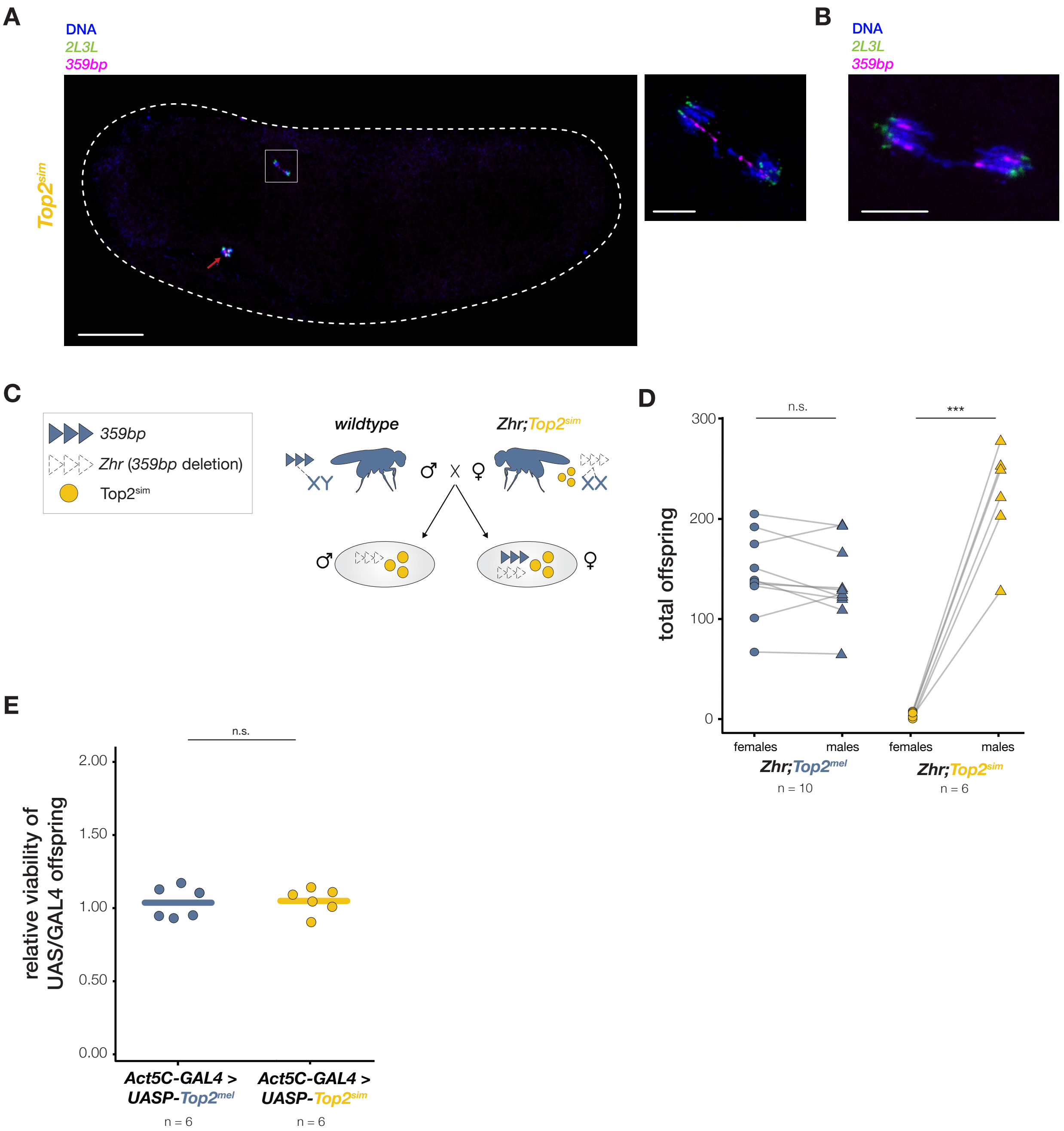




*Supplemental Figure 1*

**Fig. S1. Top2$^{sim}$-mediated lethality is restricted to the early embryo.** (**A**) Representative image of *2L3L* (green) and *359bp* (pink) segregation during the first mitotic division in a *Top2$^{sim}$*-provisioned embryo. The three residual polar bodies (arrow) are retained after female meiosis (scale bar, 50μm). Inset displays *359bp* chromatin bridge at anaphase (scale bar, 5μm). (**B**) A *Top2$^{sim}$*-provisioned embryo at the first mitosis with neither probe detected in the bridge; however, the positioning and shape of the *359bp* signal is aberrant. Specifically, the pericentromeric *359bp* array appears in the midzone rather than at the poles, and the signal indicates that the array is stretched rather than punctate. (**C**) Schematic of the cross used to assay the consequences of Top2$^{sim}$ expression in the *D. melanogaster* ovary versus maternal deposition of Top2$^{sim}$ into *D. melanogaster* embryos. Wild-type *D. melanogaster* males are crossed to transgenic *D. melanogaster* Zhr;*Top2$^{sim}$* females lacking the *X*-linked *359bp* array. These females express Top2$^{sim}$ in the ovary, but maternal Top2$^{sim}$ only encounters *359bp* derived from the paternal *X* chromosome during embryogenesis. If the incompatibility originates during oogenesis, maternal deletion of the *359bp* array would rescue the viability of all embryos. Conversely, if the incompatibility arises during embryogenesis, daughters that inherit the paternal *X*-linked *359bp* array would be inviable and *Y*-bearing sons would be viable. (**D**) Total female and male offspring from *Zhr;Top2$^{mel}$* (left) and *Zhr;Top2$^{sim}$* (right) females crossed to wild-type males. *Zhr;Top2$^{mel}$* females produce equal numbers of male and female progeny, whereas *Zhr;Top2$^{sim}$* females produce almost exclusively male progeny, indicating that the incompatibility between Top2$^{sim}$ and *359bp* arises during embryogenesis. (**E**) Relative viability of flies expressing Top2$^{mel}$ or Top2$^{sim}$ ubiquitously in the soma using the *Act5C-GAL4* driver. Viability = proportion of UAS/GAL4 progeny relative to their UAS/TM6 balancer siblings from the same cross. t-test, ***$p < 0.001$

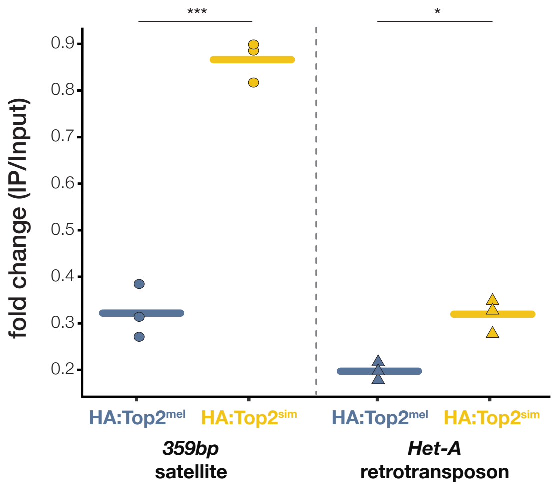


*Supplemental Figure 2*

**Fig. S2. Top2$^{sim}$ hyperaccumulates at *359bp* relative to Top2$^{mel}$**. ChIP-qPCR analysis on ovaries quantifying HA::Top2$^{mel}$ or HA::Top2$^{sim}$ enrichment at either the *359bp* satellite array or the HeT-A telomeric retrotransposon (control). Both versions of Top2 are enriched at *359bp*; however, Top2$^{sim}$ accumulates at markedly higher levels than Top2$^{mel}$. At the telomere, we observed a comparatively modest, but marginally significant, enrichment of Top2$^{sim}$ relative to Top2$^{mel}$. t-test, * $p < 0.05$, *** $p < 0.001$.

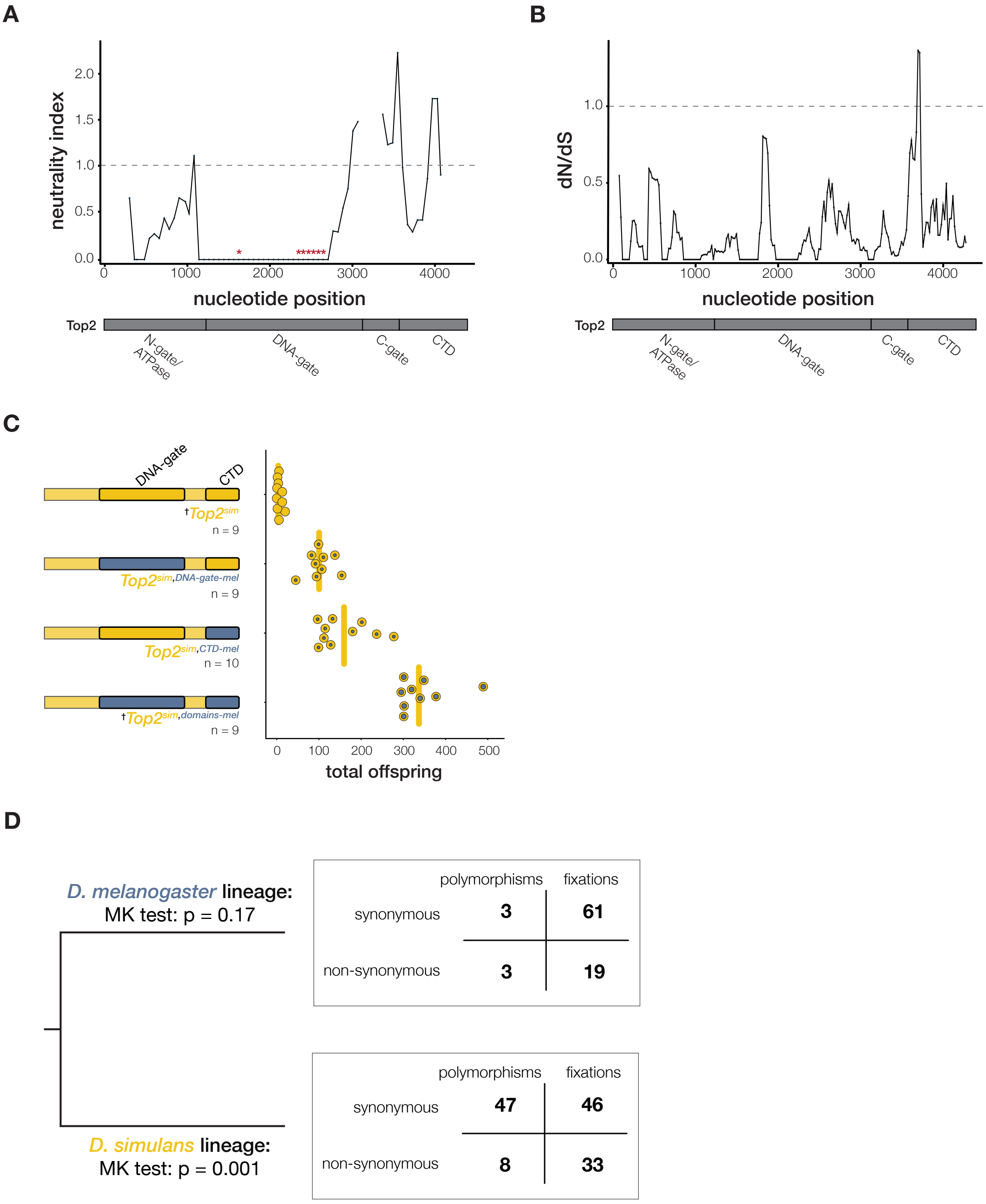


*Supplemental Figure 3*

**Fig. S3. Adaptive sequence divergence at the DNA-interacting regions of Top2 contributes to Top2 functional divergence.** (**A**) The neutrality index (NI) calculated from sliding-window McDonald-Kreitman tests across the *Top2* coding region between *D. melanogaster* and *D. simulans* (window size = 200 codons, step size = 20 codons). NI values <1 indicate an excess of nonsynonymous fixed differences, consistent with a history of positive selection. Four NI values >5 were omitted for visualization. Asterisks denote positions along Top2 with significant McDonald-Kreitman tests ($\chi^2$ test, $p < 0.05$). Major domains across Top2 are indicated below the nucleotide position. CTD = C-terminal domain. (**B**) The estimated rate of nonsynonymous substitutions relative to synonymous substitutions (dN/dS) across the *Top2* coding region between *D. melanogaster* and *D. simulans* (window size = 100 bp, step size = 20 bp). dN/dS values >1 indicate elevated amino acid divergence, consistent with a history of positive selection. Major domains across Top2 are indicated below the nucleotide position. CTD = C-terminal domain. (**C**) Total offspring from females expressing $Top2^{sim}$ or chimeric $Top2^{sim}$ transgenes. †$Top2^{sim}$ and $Top2^{sim,mel\text{-}domains}$ data replotted from Figure 2F. The adaptively evolving DNA-gate and C-terminal domains are outlined in black. CTD = C-terminal domain. t-test, ***$p < 0.001$. (**D**) Counts of Top2 synonymous and non-synonymous polymorphisms and synonymous and non-synonymous fixed sites polarized along the *D. melanogaster* and *D. simulans* lineages. MK = McDonald-Kreitman ($\chi^2$ test).

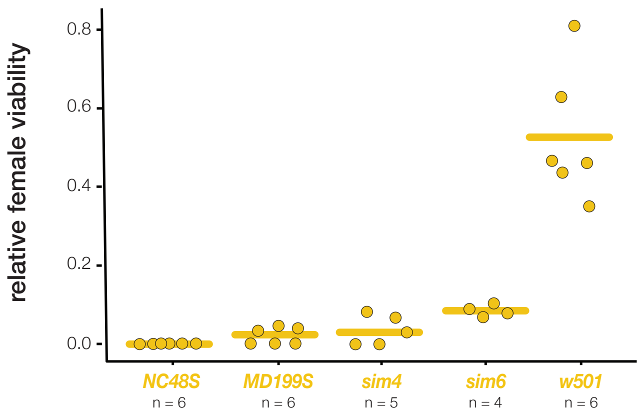


*Supplemental Figure 4*

**Fig. S4. Variation in the penetrance of maternal-effect lethality across *D. simulans* strains.** Relative viability of female offspring from crosses between males from a single *D. melanogaster* strain ($w^{1118}$) and five focal wildtype *D. simulans* strains.

```
NC48S-Top2     MENGNKGMSIEQMYQKKSQLEHILLRPDSYIGSVEFTKELMWVYDNSQNRMVQKEISFVP    60
MD199-Top2     MENGNKGMSIEQMYQKKSQLEHILLRPDSYIGSVEFTKELMWVYDNSQNRMVQKEISFVP    60
sim4-Top2      MENGNKGMSIEQMYQKKSQLEHILLRPDSYIGSVEFTKELMWVYDNSQNRMVQKEISFVP    60
sim6-Top2      MENGNKGMSIEQMYQKKSQLEHILLRPDSYIGSVEFTKELMWVYDNSQNRMVQKEISFVP    60
w501-Top2      MENGNKGMSIEQMYQKKSQLEHILLRPDSYIGSVEFTKELMWVYDNSQNRMVQKEISFVP    60
               ************************************************************

NC48S-Top2     GLYKIFDEILVNAADNKQRDKSMNTIKIDIDPERNIVSVWNNGQGIPVTMHKEQKMYVPT    120
MD199-Top2     GLYKIFDEILVNAADNKQRDKSMNTIKIDIDPERNIVSVWNNGQGIPVTMHKEQKMYVPT    120
sim4-Top2      GLYKIFDEILVNAADNKQRDKSMNTIKIDIDPERNIVSVWNNGQGIPVTMHKEQKMYVPT    120
sim6-Top2      GLYKIFDEILVNAADNKQRDKSMNTIKIDIDPERNIVSVWNNGQGIPVTMHKEQKMYVPT    120
w501-Top2      GLYKIFDEILVNAADNKQRDKSMNTIKIDIDPERNIVSVWNNGQGIPVTMHKEQKMYVPT    120
               ************************************************************

NC48S-Top2     MIFGHLLTSSNYNDDEKKVTGGRNGYGAKLCNIFSTSFTVETATKEYRKSFKQTWGNNMG    180
MD199-Top2     MIFGHLLTSSNYNDDEKKVTGGRNGYGAKLCNIFSTSFTVETATKEYRKSFKQTWGNNMG    180
sim4-Top2      MIFGHLLTSSNYNDDEKKVTGGRNGYGAKLCNIFSTSFTVETATKEYRKSFKQTWGNNMG    180
sim6-Top2      MIFGHLLTSSNYNDDEKKVTGGRNGYGAKLCNIFSTSFTVETATKEYRKSFKQTWGNNMG    180
w501-Top2      MIFGHLLTSSNYNDDEKKVTGGRNGYGAKLCNIFSTSFTVETATKEYRKSFKQTWGNNMG    180
               ************************************************************

NC48S-Top2     KASDVQIKDFNGTDYTRITFSPDLAKFKMDRLDEDIVALMSRRAYDVAASSKGVSVFLNG    240
MD199-Top2     KASDVQIKDFNGTDYTRITFSPDLAKFKMDRLDEDIVALMSRRAYDVAASSKGVSVFLNG    240
sim4-Top2      KASDVQIKDFNGTDYTRITFSPDLAKFKMDRLDEDIVALMSRRAYDVAASSKGVSVFLNG    240
sim6-Top2      KASDVQIKDFNGTDYTRITFSPDLAKFKMDRLDEDIVALMSRRAYDVAASSKGVSVFLNG    240
w501-Top2      KASDVQIKDFNGTDYTRITFSPDLAKFKMDRLDEDIVALMSRRAYDVAASSKGVSVFLNG    240
               ************************************************************

NC48S-Top2     NKLAVRNFKDYIDLHIKSTDDDSGPPIKIVHEVANERWEVACCPSDRGFQQVSFVNSIAT    300
MD199-Top2     NKLAVRNFKDYIDLHIKSTDDDSGPPIKIVHEVANERWEVACCPSDRGFQQVSFVNSIAT    300
sim4-Top2      NKLAVRNFKDYIDLHIKSTDDDSGPPIKIVHEVANERWEVACCPSDRGFQQVSFVNSIAT    300
sim6-Top2      NKLAVRNFKDYIDLHIKSTDDDSGPPIKIVHEVANERWEVACCPSDRGFQQVSFVNSIAT    300
w501-Top2      NKLAVRNFKDYIDLHIKSTDDDSGPPIKIVHEVANERWEVACCPSDRGFQQVSFVNSIAT    300
               ************************************************************

NC48S-Top2     YKGGRHVDHVVDNLIKQLLEVLKKKNKGGINIKPFQVRNHLWVFVNCLIENPTFDSQTKE    360
MD199-Top2     YKGGRHVDHVVDNLIKQLLEVLKKKNKGGINIKPFQVRNHLWVFVNCLIENPTFDSQTKE    360
sim4-Top2      YKGGRHVDHVVDNLIKQLLEVLKKKNKGGINIKPFQVRNHLWVFVNCLIENPTFDSQTKE    360
sim6-Top2      YKGGRHVDHVVDNLIKQLLEVLKKKNKGGINIKPFQVRNHLWVFVNCLIENPTFDSQTKE    360
w501-Top2      YKGGRHVDHVVDNLIKQLLEVLKKKNKGGINIKPFQVRNHLWVFVNCLIENPTFDSQTKE    360
               ************************************************************

NC48S-Top2     NMTLQQKGFGSKCTLSDKFINNMSKSGIVESVLAWAKFKAQNDIAKTGGRKSSKIKGIPK    420
MD199-Top2     NMTLQQKGFGSKCTLSDKFINNMSKSGIVESVLAWAKFKAQNDIAKTGGRKSSKIKGIPK    420
sim4-Top2      NMTLQQKGFGSKCTLSDKFINNMSKSGIVESVLAWAKFKAQNDIAKTGGRKSSKIKGIPK    420
sim6-Top2      NMTLQQKGFGSKCTLSDKFINNMSKSGIVESVLAWAKFKAQNDIAKTGGRKSSKIKGIPK    420
w501-Top2      NMTLQQKGFGSKCTLSDKFINNMSKSGIVESVLAWAKFKAQNDIAKTGGRKSSKIKGIPK    420
               ************************************************************

NC48S-Top2     LEDANEAGGKNSINCTLILTEGDSAKSLAVSGLGVIGRDLYGVFPLRGKLLNVREANYKQ    480
MD199-Top2     LEDANEAGGKNSINCTLILTEGDSAKSLAVSGLGVIGRDLYGVFPLRGKLLNVREANYKQ    480
sim4-Top2      LEDANEAGGKNSINCTLILTEGDSAKSLAVSGLGVIGRDLYGVFPLRGKLLNVREANYKQ    480
sim6-Top2      LEDANEAGGKNSINCTLILTEGDSAKSLAVSGLGVIGRDLYGVFPLRGKLLNVREANYKQ    480
w501-Top2      LEDANEAGGKNSINCTLILTEGDSAKSLAVSGLGVIGRDLYGVFPLRGKLLNVREANYKQ    480
               ************************************************************

NC48S-Top2     LSENAEVNNLCKIIGLQYKKKYLTEDDLKTLRYGKVMIMTDQDQDGSHIKGLLINFIHTN    540
MD199-Top2     LSENAEVNNLCKIIGLQYKKKYLTEDDLKTLRYGKVMIMTDQDQDGSHIKGLLINFIHTN    540
sim4-Top2      LSENAEVNNLCKIIGLQYKKKYLTEDDLKTLRYGKVMIMTDQDQDGSHIKGLLINFIHTN    540
sim6-Top2      LSENAEVNNLCKIIGLQYKKKYLTEDDLKTLRYGKVMIMTDQDQDGSHIKGLLINFIHTN    540
w501-Top2      LSENAEVNNLCKIIGLQYKKKYLTEDDLKTLRYGKVMIMTDQDQDGSHIKGLLINFIHTN    540
               ************************************************************
```

```
NC48S-Top2      WPELLRLPFLEEFITPIVKATKKNEELSFYSLPEFEEWKNDTANHHTYNIKYYKGLGTST    600
MD199-Top2      WPELLRLPFLEEFITPIVKATKKNEELSFYSLPEFEEWKNDTANHHTYNIKYYKGLGTST    600
sim4-Top2       WPELLRLPFLEEFITPIVKATKKNEELSFYSLPEFEEWKNDTANHHTYNIKYYKGLGTST    600
sim6-Top2       WPELLRLPFLEEFITPIVKATKKNEELSFYSLPEFEEWKNDTANHHTYNIKYYKGLGTST    600
w501-Top2       WPELLRLPFLEEFITPIVKATKKNEELSFYSLPEFEEWKNDTANHHTYNIKYYKGLGTST    600
                ************************************************************

NC48S-Top2      SKEAKEYFQDMQRHRILFKYDGSVDDESIIMAFSKKHIESRKVWLTNHMDEVKRRKELGL    660
MD199-Top2      SKEAKEYFQDMQRHRILFKYDGSVDDESIIMAFSKKHIESRKVWLTNHMDEVKRRKELGL    660
sim4-Top2       SKEAKEYFQDMQRHRILFKYDGSVDDESIIMAFSKKHIESRKVWLTNHMDEVKRRKELGL    660
sim6-Top2       SKEAKEYFQDMQRHRILFKYDGSVDDESIIMAFSKKHIESRKVWLTNHMDEVKRRKELGL    660
w501-Top2       SKEAKEYFQDMQRHRILFKYDGSVDDESIIMAFSKKHIESRKVWLTNHMDEVKRRKELGL    660
                ************************************************************

NC48S-Top2      PERYLYTKGTKSITYADFINLELVLFSNADNERSIPSLVDGLKPGQRKVMFTCFKRNDKR    720
MD199-Top2      PERYLYTKGTKSITYADFINLELVLFSNADNERSIPSLVDGLKPGQRKVMFTCFKRNDKR    720
sim4-Top2       PERYLYTKGTKSITYADFINLELVLFSNADNERSIPSLVDGLKPGQRKVMFTCFKRNDKR    720
sim6-Top2       PERYLYTKGTKSITYADFINLELVLFSNADNERSIPSLVDGLKPGQRKVMFTCFKRNDKR    720
w501-Top2       PERYLYTKGTKSITYADFINLELVLFSNADNERSIPSLVDGLKPGQRKVMFTCFKRNDKR    720
                ************************************************************

NC48S-Top2      EVKVAQLSGSVAEMSAYHHGEVSLQMTIVNLAQNFVGANNINLLEPRGQFGTRLTGGKDC    780
MD199-Top2      EVKVAQLSGSVAEMSAYHHGEVSLQMTIVNLAQNFVGANNINLLEPRGQFGTRLTGGKDC    780
sim4-Top2       EVKVAQLSGSVAEMSAYHHGEVSLQMTIVNLAQNFVGANNINLLEPRGQFGTRLTGGKDC    780
sim6-Top2       EVKVAQLSGSVAEMSAYHHGEVSLQMTIVNLAQNFVGANNINLLEPRGQFGTRLTGGKDC    780
w501-Top2       EVKVAQLSGSVAEMSAYHHGEVSLQMTIVNLAQNFVGANNINLLEPRGQFGTRLTGGKDC    780
                ************************************************************

NC48S-Top2      ASARYIFTLMSPLTRLIYHPLDDPLLDYQVDDGQKIEPLWYLPIIPMVLVNGAEGIGTGW    840
MD199-Top2      ASARYIFTLMSPLTRLIYHPLDDPLLDYQVDDGQKIEPLWYLPIIPMVLVNGAEGIGTGW    840
sim4-Top2       ASARYIFTLMSPLTRLIYHPLDDPLLDYQVDDGQKIEPLWYLPIIPMVLVNGAEGIGTGW    840
sim6-Top2       ASARYIFTLMSPLTRLIYHPLDDPLLDYQVDDGQKIEPLWYLPIIPMVLVNGAEGIGTGW    840
w501-Top2       ASARYIFTLMSPLTRLIYHPLDDPLLDYQVDDGQKIEPLWYLPIIPMVLVNGAEGIGTGW    840
                ************************************************************

NC48S-Top2      STKISNYNPREIMNNLRKMINGQEPVVMHPWYKNFLGRIEYVSDGRYVQTGNLQILHGNR    900
MD199-Top2      STKISNYNPREIMNNLRKMINGQEPVVMHPWYKNFLGRIEYVSDGRYVQTGNLQILHGNR    900
sim4-Top2       STKISNYNPREIMNNLRKMINGQEPVVMHPWYKNFLGRIEYVSDGRYVQTGNLQILHGNR    900
sim6-Top2       STKISNYNPREIMNNLRKMINGQEPVVMHPWYKNFLGRIEYVSDGRYVQTGNLQILHGNR    900
w501-Top2       STKISNYNPREIMNNLRKMINGQEPVVMHPWYKNFLGRIEYVSDGRYVQTGNLQILHGNR    900
                ************************************************************

NC48S-Top2      LEISELPVGVWTQNYKENVLEALSNGTEKVKAVVSEYKEYHTDTTVRFVISFAPGEFERI    960
MD199-Top2      LEISELPVGVWTQNYKENVLEALSNGTEKVKAVVSEYKEYHTDTTVRFVISFAPGEFERI    960
sim4-Top2       LEISELPVGVWTQNYKENVLEALSNGTEKVKAVVSEYKEYHTDTTVRFVISFAPGEFERI    960
sim6-Top2       LEISELPVGVWTQNYKENVLEALSNGTEKVKAVVSEYKEYHTDTTVRFVISFAPGEFERI    960
w501-Top2       LEISELPVGVWTQNYKENVLEALSNGTEKVKAVVSEYKEYHTDTTVRFVISFAPGEFERI    960
                ************************************************************

NC48S-Top2      RAEEGGFHRVFKLTTTLSTNQMHAFDQNNCLRRFPTAIDILKEFYKLRREYYARRRDFLV    1020
MD199-Top2      RAEEGGFHRVFKLTTTLSTNQMHAFDQNNCLRRFPTAIDILKEFYKLRREYYARRRDFLV    1020
sim4-Top2       RAEEGGFHRVFKLTTTLSTNQMHAFDQNNCLRRFPTAIDILKEFYKLRREYYARRRDFLV    1020
sim6-Top2       RAEEGGFHRVFKLTTTLSTNQMHAFDQNNCLRRFPTAIDILKEFYKLRREYYARRRDFLV    1020
w501-Top2       RAEEGGFHRVFKLTTTLSTNQMHAFDQNNCLRRFPTAIDILKEFYKLRREYYARRRDFLV    1020
                ************************************************************

NC48S-Top2      GQLTAQADRLSDQARFILEKCEKKLVVENKQRKAMCDELLKRGYRPDPVKEWQRRIKMED    1080
MD199-Top2      GQLTAQADRLSDQARFILEKCEKKLVVENKQRKAMCDELLKRGYRPDPVKEWQRRIKMED    1080
sim4-Top2       GQLTAQADRLSDQARFILEKCEKKLVVENKQRKAMCDELLKRGYRPDPVKEWQRRIKMED    1080
sim6-Top2       GQLTAQADRLSDQARFILEKCEKKLVVENKQRKAMCDELLKRGYRPDPVKEWQRRIKMED    1080
w501-Top2       GQLTAQADRLSDQARFILEKCEKKLVVENKQRKAMCDELLKRGYRPDPVKEWQRRIKMED    1080
                ************************************************************
```

```
NC48S-Top2      AEPADEEDDEEEEAAPSVSSKAKKEKEADPEKAFKKLTDVKKFDYLLGMSMWMLTEEKKN    1140
MD199-Top2      AEPADEEDDEEEEAAPSVSSKAKKEKEADPEKAFKKLTDVKKFDYLLGMSMWMLTEEKKN    1140
sim4-Top2       AEPADEEDDEEEEAAPSVSSKAKKEKEADPEKAFKKLTDVKKFDYLLGMSMWMLTEEKKN    1140
sim6-Top2       AEPADEEDDEEEEAAPSVSSKAKKEKEADPEKAFKKLTDVKKFDYLLGMSMWMLTEEKKN    1140
w501-Top2       AEPADEEDDEEEEAAPSVSSKAKKEKEADPEKAFKKLTDVKKFDYLLGMSMWMLTEEKKN    1140
                ************************************************************

NC48S-Top2      ELLKQRDTKLSELENLRKKTPEMLWLDDLDALESKLNEVEQKERVEEQGINLKTAKALKG    1200
MD199-Top2      ELLKQRDTKLSELENLRKKTPEMLWLDDLDALESKLNEVEQKERVEEQGINLKTAKALKG    1200
sim4-Top2       ELLKQRDTKLSELENLRKKTPEMLWLDDLDALESKLNEVEQKERVEEQGINLKTAKALKG    1200
sim6-Top2       ELLKQRDTKLSELENLRKKTPEMLWLDDLDALESKLNEVEQKERVEEQGINLKTAKALKG    1200
w501-Top2       ELLKQRDTKLSELENLRKKTPEMLWLDDLDALESKLNEVEQKERVEEQGINLKTAKALKG    1200
                ************************************************************

NC48S-Top2      QKSVSTKGRKAKSIGSGAGTLDIYPDPDGEPVEFKITEEIIKKMAAAAKVAQAAKEPKKP    1260
MD199-Top2      QKSVSTKGRKAKSIGSGAGTLDIYPDPDGEPVEFKITEEIIKKMAAAAKVAQAAKEPKKP    1260
sim4-Top2       QKSVSTKGRKAKSIGSGAGTLDIYPDPDGEPVEFKITEEIIKKMAAAAKVAQAAKEPKKP    1260
sim6-Top2       QKSVSTKGRKAKSIGSGAGTLDIYPDPDGEPVEFKITEEIIKKMAAAAKVAQAAKEPKKP    1260
w501-Top2       QKSVSTKGRKAKSIGSGAGTLDIYPDPDGEPVEFKITEEIIKKMAAAAKVAQAAKEPKKP    1260
                ************************************************************

NC48S-Top2      REPKEPKVKKEPKGKQIKADPDASGGEEVDEFDAMVEGGSKTSPKAKKAAVKKEPGEKKP    1320
MD199-Top2      REPKEPKVKKEPKGKQIKADPDASGGEEVDEFDAMVEGGSKTSPKAKKAAVKKEPGEKKP    1320
sim4-Top2       REPKEPKVKKEPKGKQIKADPDASGGEEVDEFDAMVEGGSKTSPKAKKAAVKKEPGEKKP    1320
sim6-Top2       REPKEPKVKKEPKGKQIKADPDASGGEEVDEFDAMVEGGSKTSPKAKKAAVKKEPGEKKP    1320
w501-Top2       REPKEPKVKKEPKGKQIKADPDASGGEEVDEFDAMVEGGSKTSPKAKKAAVKKEPGEKKP    1320
                ************************************************************

NC48S-Top2      RQKKENGDGLKQSKIDFSKAKAKKSDDDVEEVTPRTERPGRRQASKKIDYSSLFSDEEED    1380
MD199-Top2      RQKKENGDGLKQSKIDFSKAKAKKSDDDVEEVTPRTERPGRRQASKKIDYSSLFSDEEED    1380
sim4-Top2       RQKKENGDGLKQSKIDFSKAKAKKSDDDVEEVTPRTERPGRRQASKKIDYSSLFSDEEED    1380
sim6-Top2       RQKKENGDGLKQSKIDFSKAKAKKSDDDVEEVTPRTERPGRRQASKKIDYSSLFSDEEED    1380
w501-Top2       RQKKENGDGLKQSKIDFSKAKAKKSDDDVEEVTPRTERPGRRQASKKIDYSSLFSDEEED    1380
                ************************************************************

NC48S-Top2      GGNVGSDDDNDNASDDESPKRPGKRGREDESSGGAKKKAPPKKRRAVIESDDDDIEFDDD    1440
MD199-Top2      GGNVGSDDDNDNASDDESPKRPGKRGREDESSGGAKKKAPPKKRRAVIESDDDDIEFDDD    1440
sim4-Top2       GGNVGSDDDNDNASDDESPKRPGKRGREDESSGGAKKKAPPKKRRAVIESDDDDIEFDDD    1440
sim6-Top2       GGNVGSDDDNDNASDDESPKRPGKRGREDESSGGAKKKAPPKKRRAVIESDDDDIEFDDD    1440
w501-Top2       GGNVGSDDDNDNASDDESPKRPGKRGREDESSGGAKKKAPPKKRRAVIESDDDDIEFDDD    1440
                ************************************************************

NC48S-Top2      DSDSDFN    1447
MD199-Top2      DSDSDFN    1447
sim4-Top2       DSDSDFN    1447
sim6-Top2       DSDSDFN    1447
w501-Top2       DSDSDFN    1447
                *******
```

***Supplemental Figure 5***

**Fig. S5. No evidence of Top2 amino acid variation across a sample of *D. simulans* strains.** Alignment of Top2 protein sequences from five *D. simulans* strains (NC48S, MD199S, sim4, sim6, and w501).

**A**

***Drosophila***
n=17

| | %CDS analyzed | M7 vs. M8 log likelihood | p value |
|---|---|---|---|
| ***Top2*** | 94% | 19.85 | **<0.0001** |

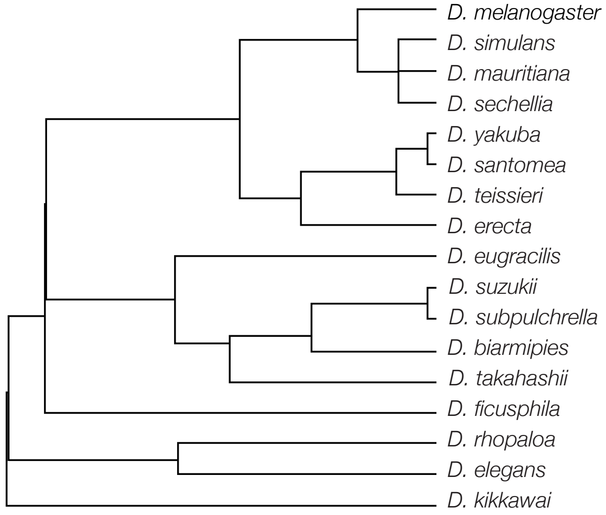


**B**

***Rodents***
n=15

| | %CDS analyzed | M7 vs. M8 log likelihood | p value |
|---|---|---|---|
| ***Top2a*** | 88% | 24.08 | **<0.0001** |
| ***Top2b*** | 100% | 0 | 1 |

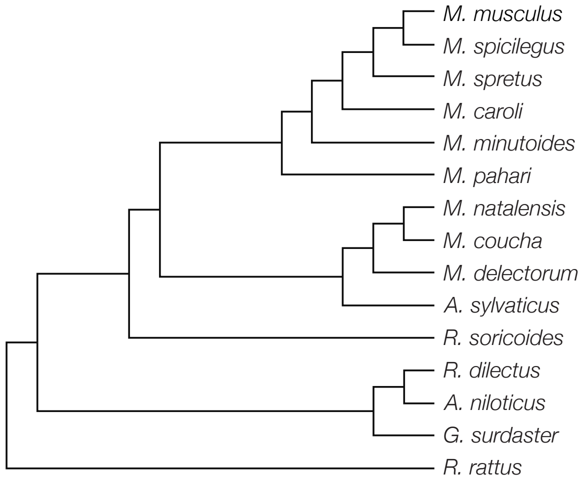


**C**

***Primates***
n=18

| | %CDS analyzed | M7 vs. M8 log likelihood | p value |
|---|---|---|---|
| ***Top2a*** | 100% | 7.79 | **0.020** |
| ***Top2b*** | 100% | 2.29 | 0.317 |

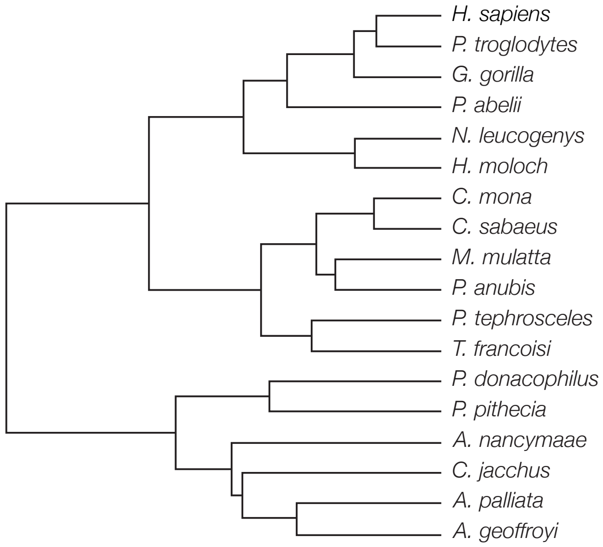


*Supplemental Figure 6*

**Fig. S6.** **Top2 evolves under positive selection across multiple eukaryotic clades.** Phylogenetic relationships of species analyzed within the (A) Drosophila, (B) rodent, and (C) primate clades and results from analyses of alternative NSsites models (Model 7 vs. Model 8) implemented in PAML. p-values were derived from a likelihood ratio test (LRT) using a chi-squared distribution.
